# Phage Commander, a software tool for rapid annotation of bacteriophage genomes using multiple programs

**DOI:** 10.1101/2020.11.11.378802

**Authors:** Matt Lazeroff, Sarah L. Harris, Philippos K. Tsourkas

## Abstract

The number of sequenced phage genomes is growing at an exponential rate. The majority of newly sequenced phage genomes are annotated by one or more of several freely-available auto-annotation programs. No program has been shown to consistently outperform the others; thus the choice of which program to use is not obvious. We present the software Phage Commander for rapid annotation of phage genomes using multiple auto-annotation programs. Phage Commander runs a phage genome sequence through nine auto-annotation programs and integrates the results within a single output. Phage Commander generates properly formatted output files for direct export to NCBI GenBank. Users can select the threshold for determining which genes should be exported. Phage Commander was benchmarked using eight high-quality phage genomes whose genes have been identified through experiments. Results show that the best results are obtained by exporting genes identified by at least two or three of the nine auto-annotation programs.

## INTRODUCTION

Each year, antibiotic-resistant bacteria cause an estimated 2.8 million infections and 35,000 deaths in the U.S. according to the U.S. Center for Disease Control (1). This figure is set to increase, as preliminary evidence suggests that part of the high mortality rate of COVID-19 could be due to opportunistic bacterial infections (2,3). Phages are attractive as an alternative to antibiotics because they are effective at lysing their host rapidly, are highly specific to their host and therefore harmless to humans and gut flora, cause few if any side effects, and co-evolve with their host, thereby reducing the chance of their host evolving resistance (4). Inconsistent early results and the development of highly successful antibiotics such as penicillin led to phages being eclipsed as treatment agents in the West. However, several recent high-profile cases of successful use of phages to resolve antibiotic-resistant infections in the US and UK have generated significant attention and renewed interest in phages as treatment agents (5–7). Phages are also becoming increasingly important in agricultural and commercial applications as treatments for infections in livestock and honeybees (8,9). They are also the source of a large number of commercial enzymes used in molecular biology (10,11).

Due to the growing interest in phages and the constantly decreasing cost of sequencing, the number of sequenced phage genomes is growing at an exponential rate (12). The sequencing of a novel phage genome is followed by annotation, which consists of 1) identifying genes, 2) identifying start codons, and 3) assigning putative function to genes (13). Genes and start codons are usually identified using one or more auto-annotation programs, such as Glimmer (14), GeneMark (15), GeneMark.hmm (16), GeneMark with Heuristics (17), GeneMarkS (18), GeneMarkS2 (19), RAST (20), BASys (21), Prodigal (22), Prokka (23), MetaGene (24), and PhANOTATE (25). Although designed for bacterial genomes (with the exception of PhANOTATE), these programs produce phage genome annotations with roughly 80-90% accuracy (12), with the added benefit of doing so rapidly. Each of these programs uses a different algorithm and produces unique results. Preliminary work has shown that no program consistently outperforms the others (12), and, thus the choice of which program to use is not obvious. Many groups combine two or three programs’ results and manually interpret their findings to achieve higher accuracy (13,26).

We have designed the program Phage Commander to facilitate running a phage genome through multiple auto-annotation programs simultaneously. Rather than trying to decide which program(s) to use, our philosophy is to combine as many programs as possible within a single user interface and integrate the results. We have thus striven to include as many auto-annotation programs as possible within a single interface. The various programs have different strengths and weaknesses; thus using multiple programs aims to increase sensitivity and specificity, provided the output is integrated appropriately.

## MATERIALS AND METHODS

Phage Commander was coded in Python 3.6+ and runs on Windows, Mac OS, and Linux. Mac and Linux users need to have Python installed on their systems and can install and run Phage Commander following the instructions on GitHub. For Windows, a standalone executable is available. Phage Commander is freely available for download, along with the source code, from Github (https://github.com/mlazeroff/PhageCommander).

Phage Commander incorporates the following nine auto-annotation programs for gene and start codon identification: Glimmer, GeneMark, GeneMark.hmm, GeneMark S, GeneMark with Heuristics, GeneMark S2, Prodigal, RAST, and MetaGene. The list of gene identification programs included in Phage Commander is given in Table 1 below. In addition to these, Phage Commander also includes the program Aragorn for tRNA identification.

**Table 1.**
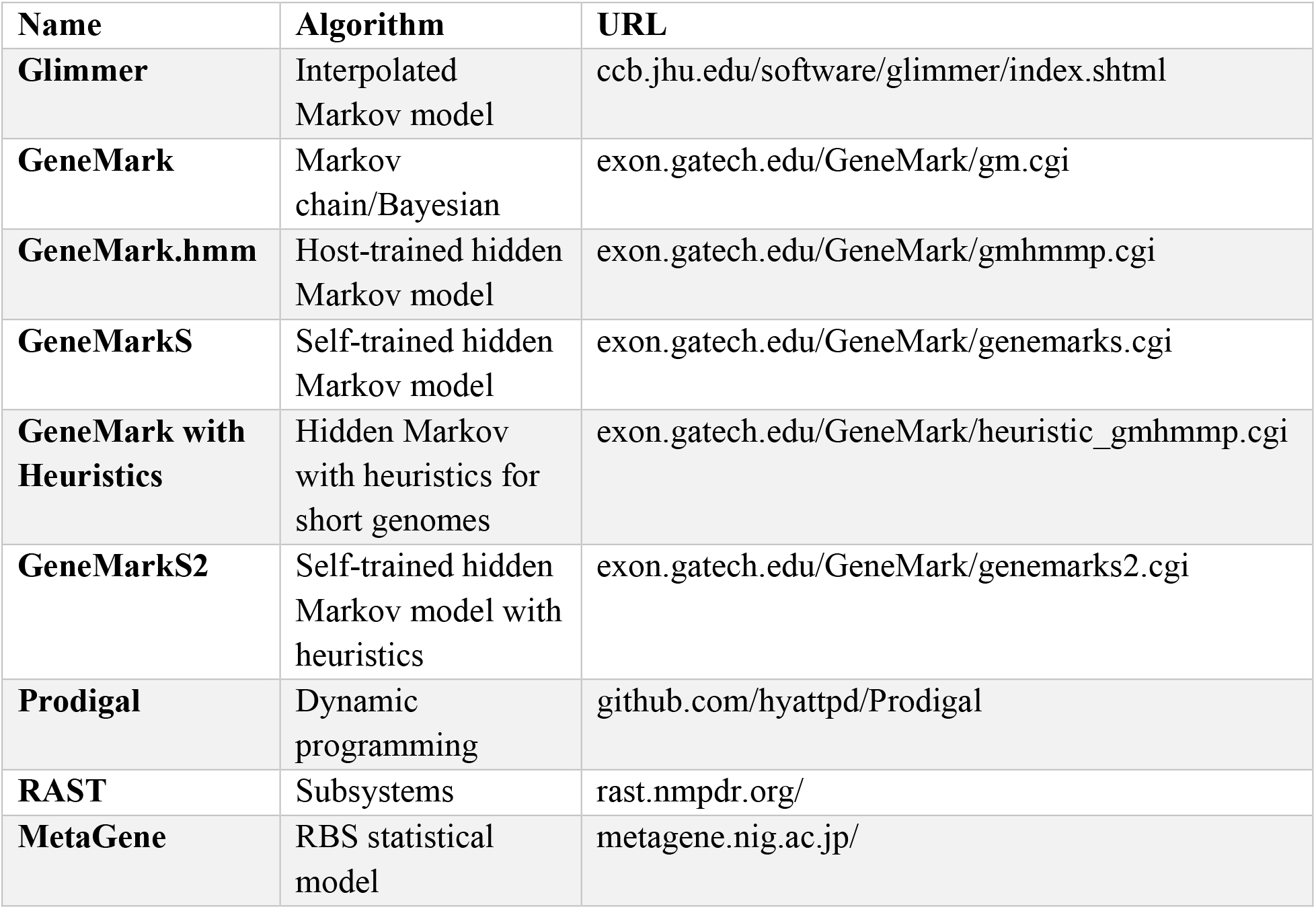
Auto-annotation programs for gene identification included in Phage Commander.

The input to Phage Commander is a phage genome sequence in fasta format. By clicking “New” in the File menu, users select the programs and fasta file to use. If using GeneMark.hmm, users need to select the bacterial host, as this is required by GeneMark.hmm. If using RAST, users should create an account on the RAST server (https://rast.nmpdr.org/rast.cgi) and enter their RAST credentials when prompted by Phage Commander.

The output of Phage Commander is a list of genes predicted by each program (start codon location, stop codon location, and length) in spreadsheet format. A sample screenshot is shown in Figure 1. Each row represents a gene, each set of four columns corresponds to one of the nine programs. For each gene, the strand, start codon, stop codon, and length are listed. Gene rows are shaded based on how many auto-annotation programs identify that particular gene, with darker shading corresponding to more program calls. The leftmost column is the number of programs that identify a gene; however which programs identify a particular gene may vary. For example, in Figure 1, gene 70 and gene 90 are both identified by three programs, but not by the same three programs. The “ALL” and “ONE” columns indicate a gene is called by all programs or only one program, respectively. The output from Aragorn is given in a separate tab.

**Figure 1.**
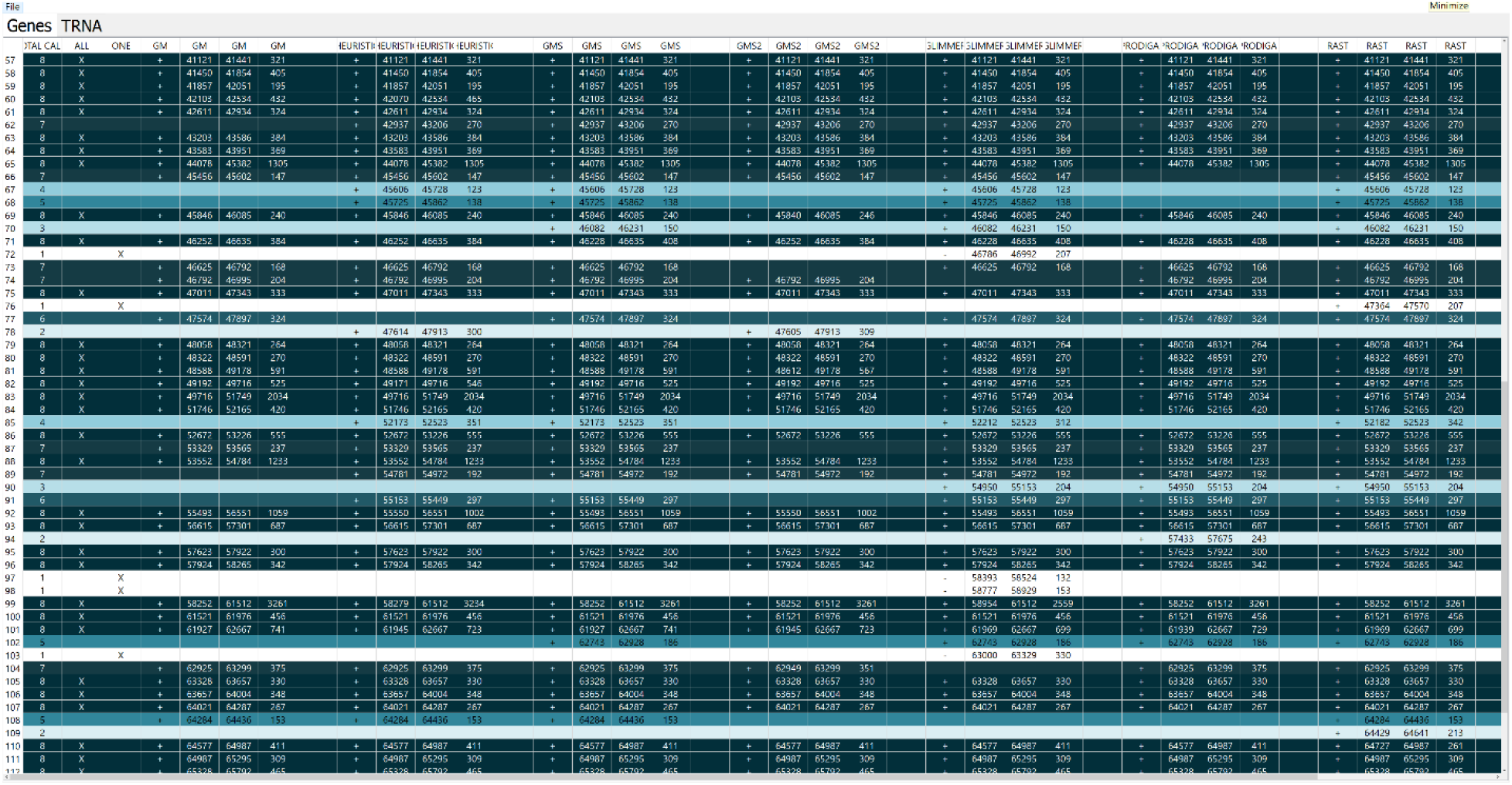
Sample Phage Commander output. Programs shown include GeneMark, GeneMark with heuristics, GeneMark S, GeneMark S2, Glimmer, Prodigal, and RAST (GeneMark.hmm and MetaGene not shown). Each gene is a row, and shaded based on how many programs identify it.

Two export options exist: Excel format (.xlsx) and GenBank format (.gb). GenBank formatted files can be directly uploaded to the NCBI GenBank genome repository. In our workflow, we export the genes predicted from Phage Commander in .gb format and read them into the DNA Master software (13), which we use to assign putative functions. The fully annotated genomes can then be again exported from DNA Master in .gb format for upload to NCBI.

When exporting the output in GenBank format, users have the ability to set the threshold for exporting genes, in terms of the number of programs that identify genes. For example, users can select to export genes identified by at least 1 program (i.e. equivalent to logical ANY/OR), genes identified by at least two programs, all the way up to genes identified by all nine programs (equivalent to logical ALL/AND). The start codons exported are those chosen by a majority rule.

## RESULTS

To benchmark Phage Commander, we searched for phage genomes whose genes are known with a high degree of certainty through experiments. We identified eight phages genomes that meet this criterion, shown in Table 2 below.

**Table 2.** Phages used in testing Phage Commander

| Name | Host | Genome length (bp) | No. of genes | GenBank accession no. |
| --- | --- | --- | --- | --- |
| **Lambda** | *E. coli* | 48,502 | 71* | NC_001416 |
| **Mu** | *E. coli* | 36,717 | 55 | NC_000929 |
| **Patience** | *M. smegmatis* | 70,506 | 110 | NC_023691 |
| **Kampy** | *M. smegmatis* | 51,378 | 89** | NC_024141 |
| **Giles** | *M. smegmatis* | 53,746 | 85*** | NC_0099933 |
| **PAK_P3** | *P. aeruginosa* | 88,097 | 165 | NC_022970 |
| **YuA** | *P. aeruginosa* | 58,663 | 79** | NC_010116 |
| **API480** | *P. larvae* | 45,026 | 77 | MK533143 |

\*2 duplicate genes not counted
\*\*Includes genes added
\*\*\*Based on the annotation of LilHazelNut

The *E. coli* phage Lambda was isolated in 1950 and is perhaps the most studied phage in existence (27). Phage Mu is another *E. coli* phage that has been extensively studied since it was isolated in the early 1960s, and whose genome has been carefully annotated (28). Phage Patience is a somewhat atypical *Mycobacterium smegmatis* phage whose genome has been extensively studied through RNA-Seq transcriptomics and mass spectrometry (29). Phage Kampy is a cluster A4 *M. smegmatis* phage that has also been studied through transcriptomics and mass spectrometry (30). Phage Giles is a cluster Q *M. smegmatis* phage that has been studied through mass spectrometry and whose protein interactome has been mapped out (31). Phage PAK_P3 is a *Pseudomonas aeruginosa* phage that has been studied through transcriptomics experiments (32). Phage YuA is a *P. aeruginosa* phage that has been studied through mass spectrometry and other experiments (33). Phage API480 is a phage that infects the bacterium *Paenibacillus larvae,* a pathogen of the honeybee *Apis mellifera,* and has been the subject of transcriptomics experiments (34). The published annotations of these eight phage genomes in NCBI GenBank are considered highly trustworthy and were used as references to benchmark the performance of Phage Commander. A gene that is detected by transcriptomics or mass spectrometry is likely to be real, so we did not remove any genes from the reference annotations. However, even transcriptomics might miss some genes that are rarely expressed. We thus inspected each genome carefully and added one gene to the reference genome of Kampy and two genes to YuA that were identified by all 9 programs, had many statistically significant homologs, and filled a coding gap. For phage Giles, we used the annotation of the closely related phage LilHazelnut (99.99% nucleotide identity), whose annotation is more recent and appears more complete (35). Lambda has two cases of two genes within the same reading frame, differing only by their start (genes nu3 and D, and genes S and R); we thus counted the duplicate genes as a single gene (i.e. we counted nu3 and D as one gene, and S and R as one gene), because no program has the ability to identify two genes within the same reading frame.

Each genome was run through Phage Commander. Genes were identified by requiring 1-9 programs to identify a gene, and the results were compared with the reference annotations in Table 2. Figures 2 and 3 show the performance of Phage Commander compared with the reference annotations. A false positive is a gene that is identified by one or more programs but that is not present in the reference annotation. A false negative is a gene present in the reference annotation but not detected by the programs.

**Figure 2.**
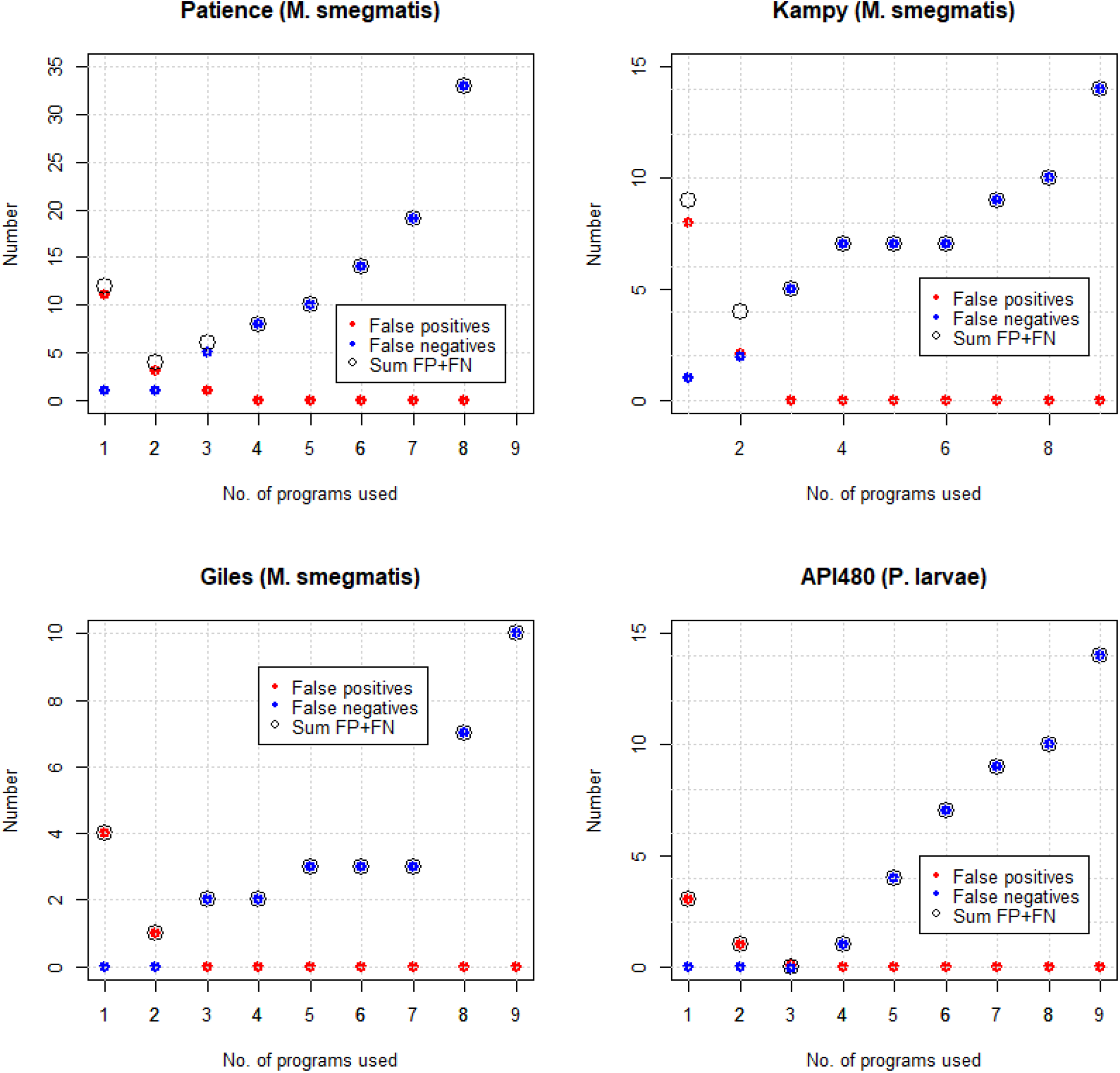
False positives, false negatives, and their sum, based on the number of programs used to export genes in Phage Commander for phages Patience, Kampy, Giles, and API480.

**Figure 3.**
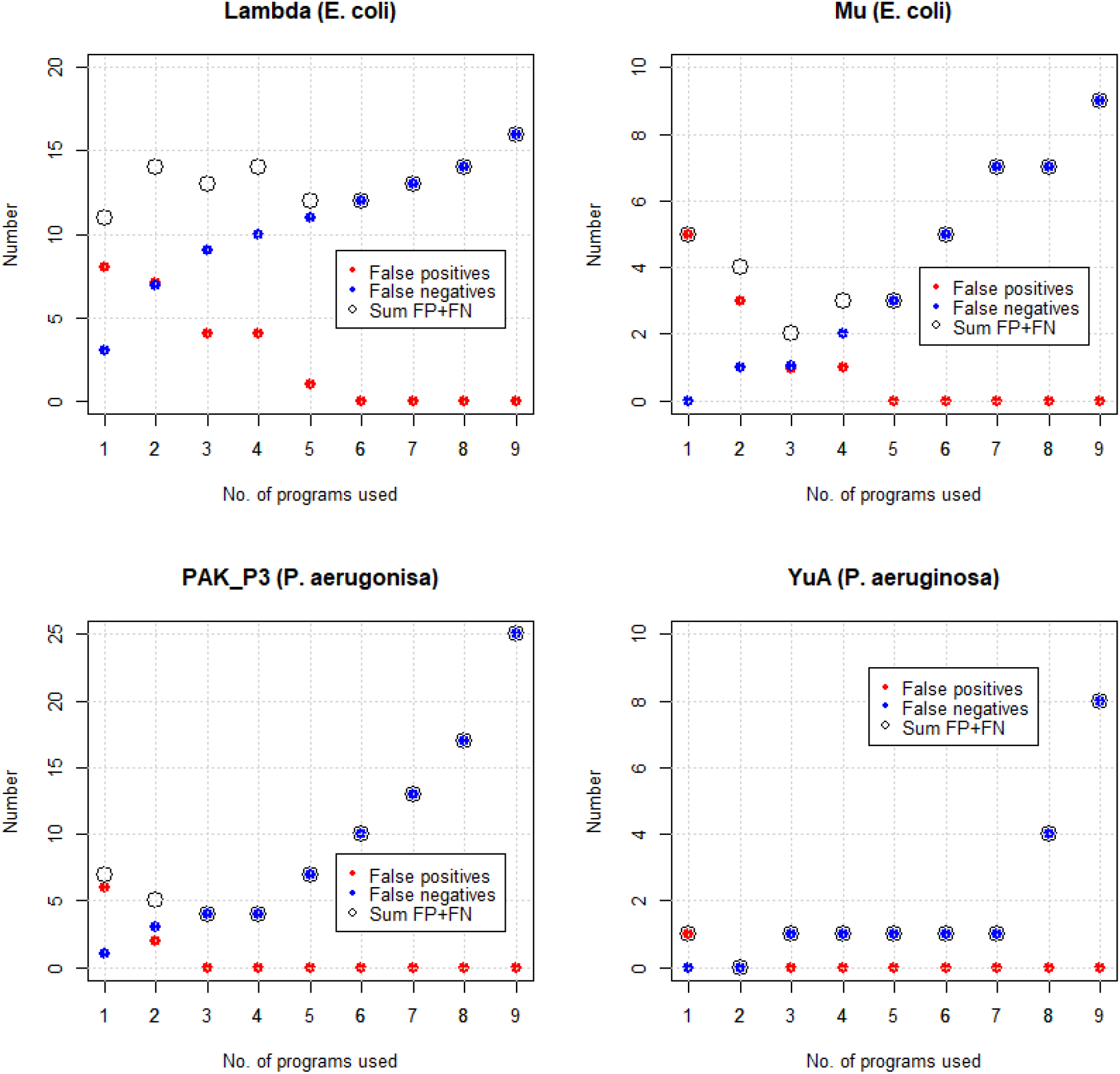
False positives, false negatives, and their sum, based on the number of programs used to export genes in Phage Commander for phages Lambda, Mu, PAK_P3 and YuA.

These results show that the sum of false positives and false negatives is usually a minimum when exporting genes called by at least two programs (Patience, Kampy, Giles, YuA), or genes called by at least three programs (Mu, PAK_P3, API480). The results for Lambda are somewhat anomalous due to the high number of false negatives. Thus, either of the “at least 2 programs” or “at least 3 programs” settings offer the best tradeoff between sensitivity and specificity, with the “at least 2 programs” setting favoring sensitivity (fewer false negatives), and the “at least 3 programs” setting favoring specificity (fewer false positives). Exporting genes called by at least one program (i.e. all genes identified) will produce a large number of false positives, although it will generate few false negatives. This setting should thus only be used if it is desired to not miss any genes without concern to the presence of false positives in the annotation. Exporting genes only called by more than 4 programs is in the opinion of the authors too stringent and will result in an unacceptably high number of false negatives (typically between 3 and 10). There is also variability between phages, with Lambda having the largest number of false positives and false negatives, and YuA the fewest, even when adjusted for genome length.

Phage Commander was developed to simplify and accelerate the manual curation method of phage genomes developed in the Tsourkas lab (12). A major component of the manual curation method is how many programs (out of nine) identify a candidate gene when making a gene identification decision (in addition to additional information such as homology, operons, overlap, ribosome binding score, and synteny) (12). Prior to the development of Phage Commander, phage genomes had to be run through each auto-annotation program separately and the output of each program was manually added into a single spreadsheet. Phage Commander was developed to accelerate the annotation process by automating running the phage genome through the auto-annotation programs and outputting the results into a single spreadsheet automatically. By our estimate, using Phage Commander accelerates our workflow by ~33%. Of interest is the relative performance of each of the nine programs used and the manual curation method, which integrates all the programs. In the original publication presenting the method, we benchmarked the method on phages Lambda and Patience (12). In Figures 4 and 5, we compare the performance of the nine programs and the manual curation method for the phages in this study.

**Figure 4.**
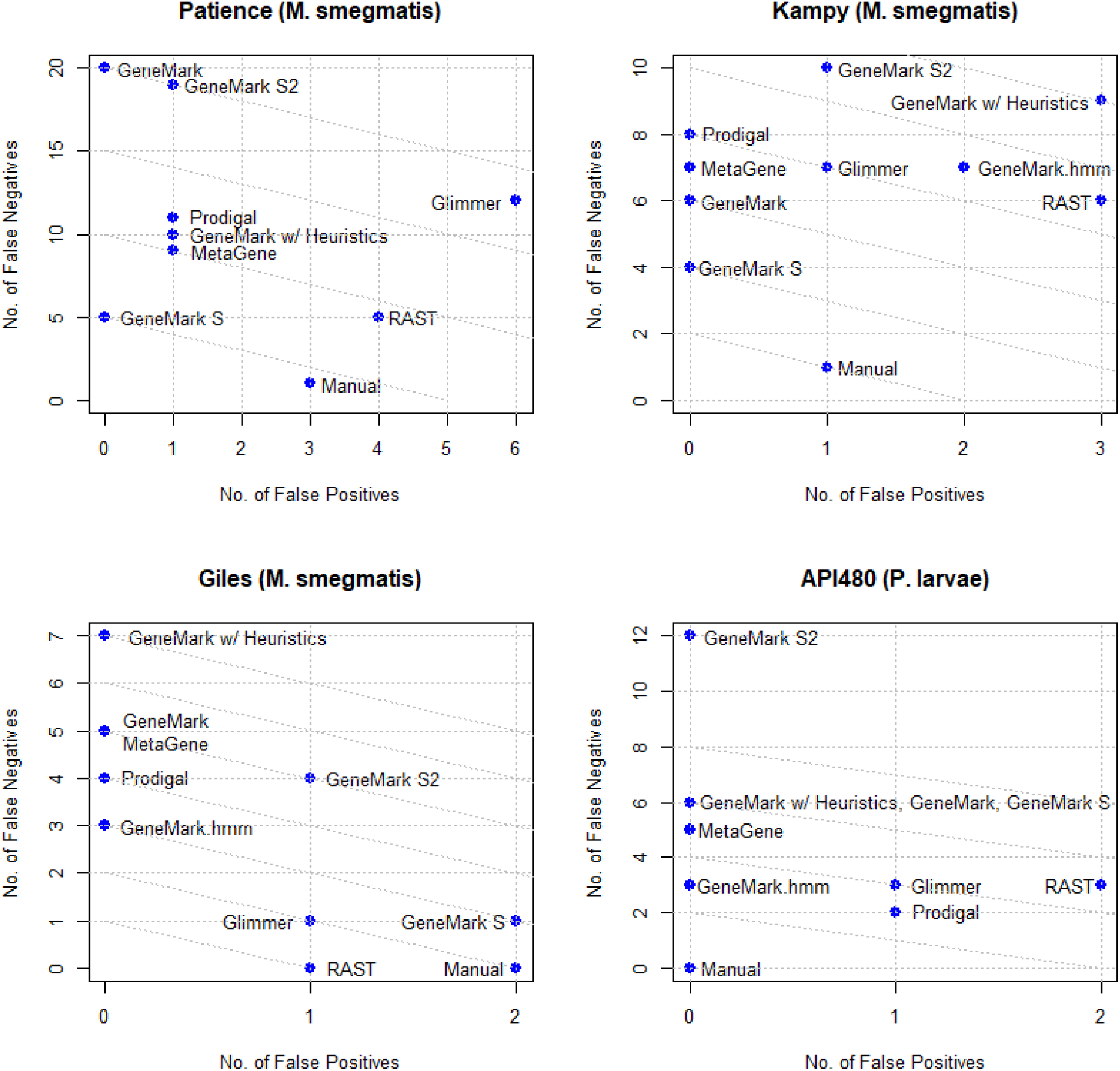
Plots of false positives vs. false negatives for Patience, Kampy, Giles, and API 480. Diagonal lines represent equal numbers of false positives and false negatives.

**Figure 5.**
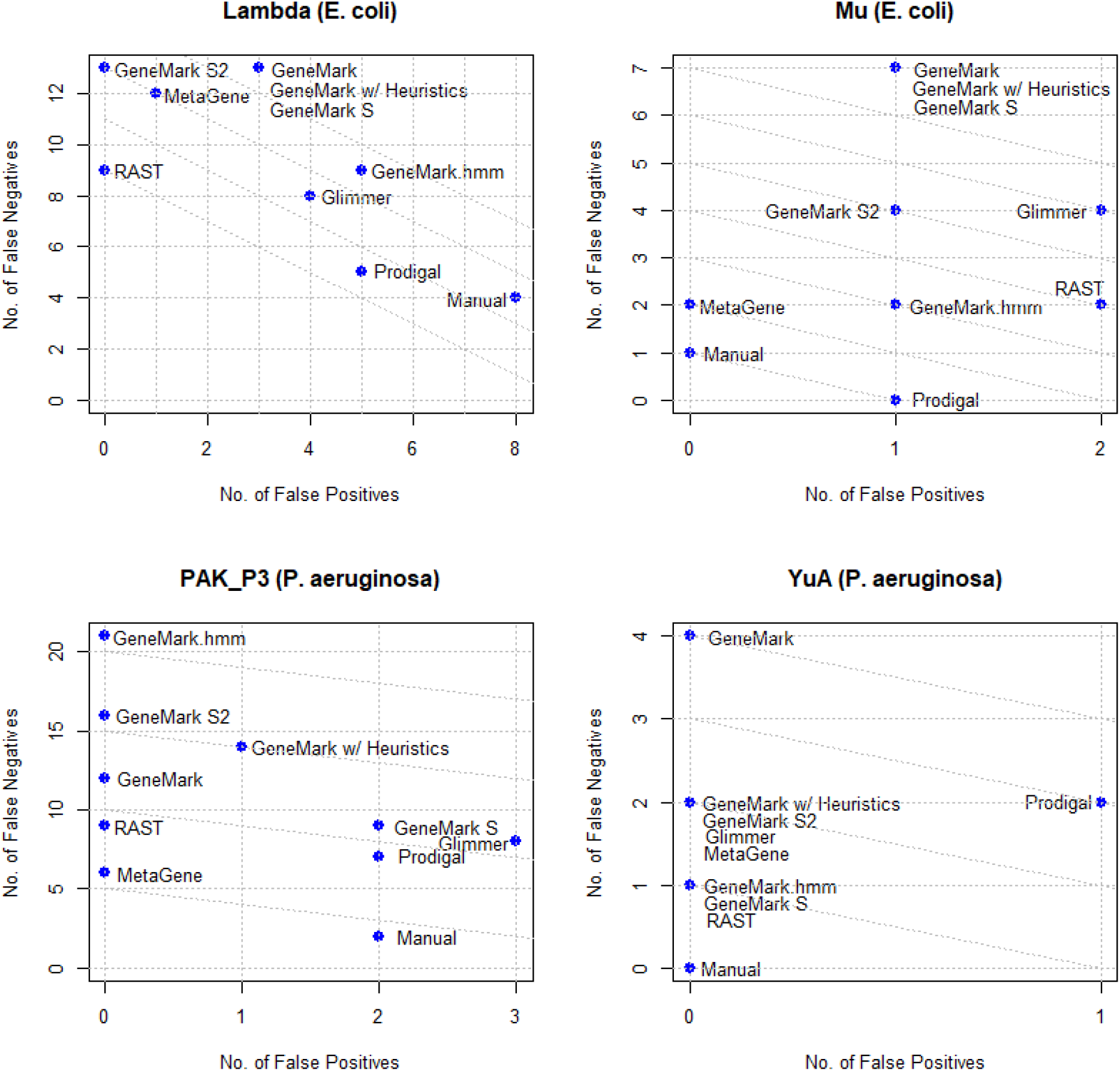
Plots of false positives vs. false negatives for Lambda, Mu, PAK_P3, and YuA. Diagonal lines represent equal numbers of false positives and false negatives.

Manual curation produces the best results (fewest false positives and false negatives) in all phages except Lambda and Giles, and tied with Prodigal in phage Mu. Also of note is that manual curation produced the fewest false negatives in all phages except Mu. Manual curation has zero false negatives for phages Giles, API480 and YuA, and achieved perfect results (zero false positives and false negatives) for phages API480 and YuA.

Of the programs, RAST appears to be the most consistently sensitive (fewest false negatives), while GeneMark, GeneMark with heuristics, and GeneMark S2 are the least sensitive on average (highest false negatives). MetaGene consistently generates the fewest false positives (never more than one, and usually zero). Of the GeneMark suite, GeneMark S and GeneMark.hmm appear to have the best performance. GeneMark.hmm is host-trained and cannot be used when the host is not known (e.g. metagenomics) or when the host is unusual. For example, GeneMark.hmm was not used for phage Patience, as this phage is an atypical *M smegmatis* phage that appears to transitioning from another host to*M. smegmatis,* and thus GeneMark.hmm gave highly anomalous results for this phage.

## DISCUSSION

Phage Commander is a software tool designed to accelerate time-consuming manual curation methods for phage genomes. It incorporates ten auto-annotation programs (nine for protein genes and one for tRNA genes) that are widely used in phage genomics within a single interface and then summarizes the results within a single spreadshet to clearly convey and visualize results. Users have the ability to select which auto-annotation programs to use and which genes to export based on the number of programs identifying a gene. Phage Commander has the ability to export files in GenBank format (.gb) for direct deposition to NCBI GenBank, or for further processing through DNA Master.

An important feature of Phage Commander is the ability for users to set the threshold for exporting genes in terms of the number of programs. Users can export genes on a sliding scale, starting from exporting all genes called by at least one program (logical ANY/OR) to only exporting genes called by all nine programs (logical ALL/AND). Phage Commander was tested on eight high quality annotated phage genomes the majority of whose genes are experimentally verified. Results show that the optimal settings are to export genes called by at least two or at least three programs. Exporting genes called by one program resulted in few false negatives (typically 0 or 1), but a high number of false positives (typically 5 to 10). Exporting genes called by more than four programs produced an unacceptably high number of false negatives (typically 3 to 10).

Of the nine programs used, results with the eight phage genomes used in this study showed RAST to be the most sensitive (fewest false negatives on average), and MetaGene the most specific (fewest false positives on average). Manual curation, which relies on integrating the output of all nine programs via Phage Commander, produced the best results (fewest combined false positives and false negatives on average) for six of the eight phages tested, showing that combining the output from multiple programs, in combination with additional information, reduces the number of false positives and false negaticves.

Future directions include adding more auto-annotation programs (e.g. PHANOTATE, BASys), and homology search results (e.g. BLAST, CD-Search, HMMer, etc.) so as to include putative function information. An additional feature to be added would be including settings for which start codons to use for exported genes. The current version exports the start codons chosen by majority rule by default, but it in future versions it will be possible to use alternative settings (e.g. choosing the start codons chosen by a specific programs). Additionally, we plan to integrate the results not just using simple logical rules (ANY, ALL), but by developing a machine learning algorithm to do so.

## CONCLUSION

We present the program Phage Commander for rapid annotation of phage genomes using multiple auto-annotation programs. By combining different programs, Phage Commander achieves better gene detection than any single auto-annotation program. Phage Commander is freely available for download from GitHub and runs on Windows, Mac and Linux.

## ACKNOWLEDGMENTS

PKT and SLH wish to acknowledge support from the UNLV Faculty Opportunity Award program for funding.

## AUTHOR CONFIRMATION STATEMENT(S)

M.L. developed the software. S.L.H. secured funding for the project and edited the paper. P.K.T designed the project, secured funding, and wrote the paper. All co-authors have reviewed and approved of the manuscript for submission. The manuscript has been submitted solely to PHAGE and is not published, in press, or submitted elsewhere.

## REFERENCES

1. Antibiotic/Antimicrobial Resistance (AR/AMR) https://www.cdc.gov/drugresistance/index.html. Access date: 10/9/2020.

2. Cevik M, Bamford C, Ho E. COVID-19 pandemic – A focused review for clinicians. Clin. Microbiol Infect 2020;7:842–847.

3. Zhou F, Yu T, Du R, et al. Clinical course and risk factors for mortality of adult inpatients with COVID-19 in Wuhan, China: A retrospective cohort study. Lancet 2020;395:1054–1062.

4. Altamirano G, Barr JJ. Phage therapy in the postantibiotic era. Clin Microbiol Rev 2019;32:143–216.

5. Schooley RT, Biswas B, Gill JJ, et al. Development and use of personalized bacteriophage-based therapeutic cocktails to treat a patient with a disseminated resistant Acinetobacter baumannii infection. Antimicrob Agents Chemother 2017;61:e00954–17.

6. Chan BK, Turner PE, Kim S, et al. Phage treatment of an aortic graft infected with Pseudomonas aeruginosa. Evol Med Pub Health 2018;1:60–66.

7. Dedrick RM, Guerrero-Bustamante CA, Garlena RA, et al. Engineered bacteriophages for treatment of a patient with a disseminated drug-resistant Mycobacterium abscessus. Nat Med 2019;25:730–733.

8. Cooper, IR. A review of current methods using bacteriophages in live animals, food and animal products intended for human consumption. J Microbiol Meth 2016;130:38–47.

9. Brady TS, Merrill BD, Hilton JA, et al. Bacteriophages as an alternative to conventional antibiotic use for the prevention or treatment of Paenibacillus larvae in honeybee hives. J Invertebr Pathol. 2017;150:94–100.

10. Grose, JH, Casjens SR. Understanding the enormous diversity of bacteriophages: the tailed phages that infect the bacterial family Enterobacteriacae. Virol 2014;468-470:421–443.

11. Kropinski AM, Clokie MRJ. Introduction. In: Clokie MRJ, Kropinski AM; eds. Bacteriophages: Methods and Protocols, Volume 2: Molecular and Applied Aspects. Valley Stream, NY: Humana Press; 2019: xiii–xxii.

12. Salisbury A, Tsourkas PK. A Method for Improving the Accuracy and Efficiency of Bacteriophage Genome Annotation. Int J Mol Sci. 2019;20(14):3391.

13. Pope WH, Jacobs-Sera, D. Annotation of bacteriophage genome sequences using DNA Master: An overview. In: Clokie MRJ, Kropinski AM; eds. Bacteriophages: Methods and Protocols, Volume 3: Molecular and Applied Aspects. Valley Stream, NY: Humana Press; 2018: 217–229.

14. Delcher AL, Harmon D, Kasif S, et al. Improved microbial gene identification with GLIMMER. Nucl Acid Res 1999;27(23):4636–4641.

15. Borodovsky M, McIninch J. GeneMark: Parallel gene recognition for both DNA strands. Comput Chem 1993;17:123–133.

16. Lukashin AV, Borodovsky M. GeneMark.hmm: New solutions for gene finding. Nucl Acids Res 1998;26(4):1107–1115.

17. Besemer J, Borodovsky M. Heuristic approach to deriving models for gene finding. Nucl Acids Res 1999;27:3911–3920.

18. Besemer J, Lomsadze A, Borodovsky M. GeneMarkS: A self-training method for gene starts in microbial genomes. Implications for finding sequence motifs in regulatory regions. Nucl Acids Res 2001;29:2607–2618.

19. Lomsadze A, Gemayel K, Tang S, et al. Modeling leaderless transcription and atypical gene results in more accurate gene prediction in prokaryotes. Genome Res 2018;20:1079–1089.

20. Aziz RK, Bartels D, Best AA, et al. The RAST Server: Rapid Annotations using Subsystems Technology. BMC Genomics 2008:8(9):75.

21. Van Domselaar GH, Stothard P, Shrivastava S, et al. BASys: a web server for automated bacterial genome annotation. Nucl Acid Res 2005;33:W455–9.

22. Hyatt D, Chen GW, LoCascio PF, et al. Prodigal: Prokaryotic gene recognition and translation initiation site identification. BMC Bioinform 2010;11:119.

23. Seemann T. Prokka: rapid prokaryotic genome annotation. Bioinformatics 2014;30:2068–9.

24. Noguchi H, Park J, Takagi T. MetaGene: prokaryotic gene finding from environmental genome shotgun sequences. Nucl Acids Res 2006;34(19):5623–30.

25. McNair K, Zhou C, Dinsdale EA, et al. PHANOTATE: A Novel Approach to Gene Identification in Phage Genomes. Bioinformatics 2019;btz265.

26. Philipson CW, Voegtly LJ, Lueder MR, Long KA, Rice GK, Frey KG, Biswas B, Cer RZ, Hamilton T, Bishop-Lilly KA. Characterizing Phage Genomes for Therapeutic Applications. Viruses. 2018 Apr 10;10(4):188. doi: 10.3390/v10040188. PMID: 29642590; PMCID: PMC5923482.

27. Casjens SR, Hendrix RW. Bacteriophage lambda: Early pioneer and still relevant. Virology. 2015;479-480:310–330.

28. Morgan GJ, Hatfull GF, Casjens S, Hendrix RW. Bacteriophage Mu genome sequence: analysis and comparison with Mu-like prophages in Haemophilus, Neisseria and Deinococcus. J Mol Biol. 2002;317(3):337–359.

29. Pope WH, Jacobs-Sera D, Russell DA, et al. Genomics and proteomics of mycobacteriophage patience, an accidental tourist in the Mycobacterium neighborhood. mBio. 2014;5(6):e02145.

30. Halleran A, Clamons S, Saha M. Transcriptomic Characterization of an Infection of Mycobacterium smegmatis by the Cluster A4 Mycobacteriophage Kampy. PLoS One. 2015;10(10):e0141100.

31. Mehla J, Dedrick RM, Caufield JH, et al. The Protein Interactome of Mycobacteriophage Giles Predicts Functions for Unknown Proteins. J Bacteriol. 2015;197(15):2508–2516.

32. Blasdel BG, Chevallereau A, Monot M, et al. Comparative transcriptomics analyses reveal the conservation of an ancestral infectious strategy in two bacteriophage genera. ISME J. 2017;11(9):1988–1996.

33. Ceyssens PJ, Mesyanzhinov V, Sykilinda N, et al. The genome and structural proteome of YuA, a new Pseudomonas aeruginosa phage resembling M6. J Bacteriol. 2008;190(4):1429–1435.

34. Ribeiro HG, Melo LDR, Oliveira H, et al. Characterization of a new podovirus infecting Paenibacillus larvae. Sci Rep. 2019;9(1):20355.

35. Shanks RA, Hazel AN, Jones WH, Segura-Totten M. Genome Sequence of Mycobacterium Phage LilHazelnut. Microbiol Resour Announc. 2019;8(19):e00431–19.

